# Nanoscale spatial dependence of avidity in an IgG1 antibody

**DOI:** 10.1101/632323

**Authors:** Agnieszka Jendroszek, Magnus Kjaergaard

## Abstract

Antibodies are secreted proteins that are crucial to recognition of pathogens by the immune system and are also efficient pharmaceuticals. The affinity and specificity of target recognition can increase remarkably through avidity effects, when the antibody can bind a multivalent antigen through more than one epitope simultaneously. A key goal of antibody engineering is thus to optimize avidity, but little is known about the nanoscale spatial dependence of avidity in antibodies. Here, we develop a set of anti-parallel coiled-coils spanning from 8-21 nm and validate their structure using biophysical techniques. We use the coiled-coils to control the spacing between two epitopes, and measure how antigen spacing affects the stability of the bivalent antibody:antigen complex. We find a maximal avidity enhancement at a spacing of 14 nm, but only see a ∼2-fold variation of avidity in the range from 8-21 nm. In contrast to recent studies, we find the avidity to be relatively insensitive to epitope spacing near the avidity maximum as long as it is within the spatial tolerance of the antibody. The coiled-coil systems developed here may prove a useful protein nanocaliper for profiling the spatial tolerance and avidity profile of bispecific antibodies.

## Introduction

Antibodies are proteins secreted by the immune system that detect and neutralize foreign molecules. Antibodies contain variable regions that can be combined to generate an abundance of different sequences, which means that they can recognize almost any other molecule. This ability has made antibodies a valuable research tool in biology and recently a successful class of pharmaceuticals.^1^ The most abundant class of antibodies (IgG) may bind two identical epitopes simultaneously if the antigens are multivalent or are closely spaced on a surface. In such cases, the combined affinity may be much greater than that for a monovalent interaction, which is known as avidity.^2^ Currently, we know little about how avidity in antibodies depends on the spatial arrangement of epitopes, which prevents the rational use of avidity in antibody engineering. Optimization of avidity and multivalency will be particularly important to bispecific antibodies, where engagement of several epitopes is essential to the therapeutic effect.

Antibodies are flexible, multidomain proteins allowing their structure to adopt to many different antigens. A common antibody scaffold is shared among the whole class of molecules (Fig. 1A). For IgG antibodies, the binding regions are located at the variable tips of two Fab moieties, and that are connected to the Fc moiety via flexible hinges. The hinges vary between IgG sub-types, and contain interchain disulfide bonds that connect the two Fab moieties. The flexible hinges allow considerable internal freedom between the Fc- and the Fab-moieties. In the unbound state, antibodies thus occupy an ensemble of different conformations, where the intra-molecular dimensions reach up to 19 nm.^3,4^ Full-length antibodies rarely crystallize, likely because domain orientation is still dynamic in most complexes.^5^ However, rare crystal structures of full-length antibodies suggested that a fully extended hinge (Fig. 1A) would allow a distance up to 9 nm between the binding site and the first disulfide bond in the hinge. This suggests a total reach for bivalent binding of antibodies of ∼18 nm,^6^ which is mirrored by the ensemble occupied in the free state.

**Figure 1.**
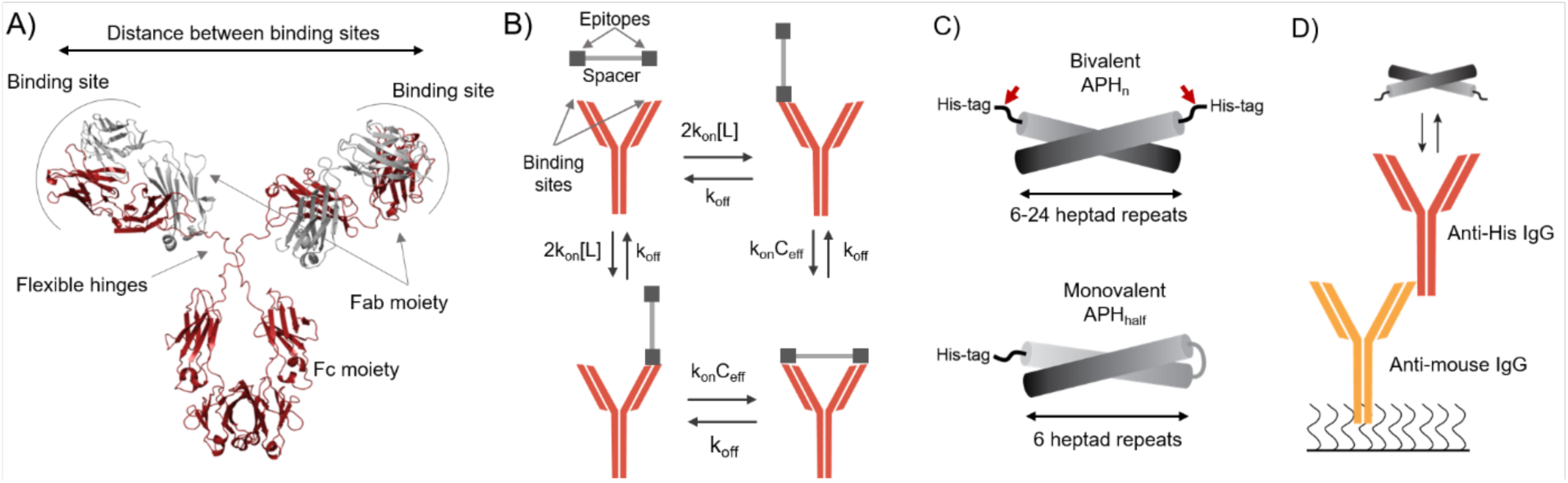
Experimental setup for testing the spatial dependence of avidity in an IgG1 antibody. (A) Domain architecture of an IgG antibody. The flexible hinges allow the two Fab moieties to move independently from the Fc module, and determine the distance tolerated between the two epitopes. (B) Schematic illustration of a bivalent binding between an antibody and an antigen with two epitopes separated by a rigid nanocaliper. The initial association is followed by a an intra-complex ring closing reaction. (C) The design of a bi- and monovalent antigens used to probe the spatial dependence of avidity. The bivalent antigen is composed of two monomers each containing an α-helical segment connected to an epitope (His-tag). Helices are composed of six to twenty four heptad repeats and are capable of formation of anti-parallel coiled-coils, which serve as rigid spacers between two epitopes. A thrombin site was introduced between the His-tag and the helix (red arrow) to allow removal of the tag prior to SAXS, DLS and SR-CD experiments. The monovalent antigen is composed of a single His-tag connected to six heptad repeats followed by a flexible loop and six matching heptad repeats, which forms an intramolecular anti-parallel coiled coil. (D) The SPR setup used for avidity measurements includes an anti-mouse antibody targeting the Fc region of the mouse IgG covalently bound to the SPR chip. In each cycle monoclonal mouse anti-His-tag Ab is captured via interaction with anti-mouse antibody. This allows oriented capture without affecting antibody flexibility and antigen binding.

Avidity affects both the association and dissociation step of a binding reaction. The association rate increases as the antibody can bind to several sites, which simply increase the association rate-constant by the multiplicity of the reaction. Following the initial association, the other Fab moiety can bind in an intra-molecular reaction called ring-closing (Fig. 1B). Ring-closing occurs intra-molecularly and is thus independent of the concentration. Instead, it depends on the structure of the antibody and antigen, which together define an effective concentration.^7,8^ Dissociation from a bivalent target requires simultaneous release of both Fab moieties, and thus depends on the ring-closing equilibrium and the effective concentration. In principle, the avidity of a bivalent interaction could be predicted from the effective concentration of ring-closing.^9^ Effective concentrations and avidity has previously been studied using either model systems^7,8^ or theoretical models.^10–12^ Neither can fully mimic the complex energy landscape of a dynamic antibody, where bivalent binding will be accompanied by changes in the structural ensemble of the antibody.

The conserved structure of the antibody scaffold suggests that the spatial dependence of avidity may by understood for one antibody and extrapolated to other antibodies in its class. The distance between the epitopes is likely to be one of the principal determinant of avidity. In antibody engineering, it is thus critical to know at which epitope spacing the avidity is the greatest. Similarly, it is critical to determine how large an epitope separation can be tolerated by the antibody, and how strongly avidity depends on the epitope spacing. To answer these questions, we wanted to vary the distance between epitopes in a controlled manner. Inspired by recent successes of DNA nanocalipers^13^ and protein origami,^14,15^ we set out to design a set of coiled-coils that could act as nanocalipers between epitopes.

Coiled-coils form rigid extended structures with lengths that depend on the number of heptad repeats. We decided to use anti-parallel coiled-coils, as a linear epitope placed at the end of the helix will result in two identical epitopes at either end (Fig 1C). The coiled-coils were based on a six-repeat coiled-coil (APH_6_) designed previously,^16^ and constructed by concatenation of the APH_6_ sequence up to 24 heptads in length with additional modifications to prevent out-of-phase dimerization (sequences in the SI). We named the variant with n heptad repeats APH_n_. We also generated a monovalent coiled-coil (APH_half_) with an identical epitope-presentation by connecting two sequential APH_6_ sequences with a PGSGSGP-linker to allow formation of an intra-molecular coiled-coil (Fig. 1C). The proteins were expressed in inclusion bodies, and refolded in 3 or 4 steps before purification. The refolding efficiency decreased with increasing length, and for the longest variants was under 5%, thus preventing the series from being extended further.

The spatial tolerance is determined by the antibody scaffold, and therefore we could choose a convenient epitope as a model system. We used an N-terminal 6xHis-tag connected to the coiled-coil by a short flexible loop (GSS). The His-tag is recognized by several commercial antibodies and have an affinity that is sensitive to pH near the pK_a_ of histidine. As the antibody structure is independent of pH from 5.5-9,^17^ this allowed us to tune the monovalent affinity without changing the epitope. The antibody can be uniformly captured on SPR chips using antibodies directed against the Fc moiety of murine IgG antibodies (Fig. 1D). The capture system only recognized the Fc moiety and thus do not affect the two Fab moieties that interact with the antigen.

We characterized the structure of the APH variants by a range of biophysical techniques to test whether they form the desired structures. To isolate the pure coiled-coil before characterization, we used variants with a thrombin cleavable His-tag that only leaves three residues (GSH) before the beginning of the coiled-coil. First, we studied the variants by synchrotron-radiation CD to test whether the proteins form coiled-coils. All CD spectra are characteristic of α-helical proteins (Fig. 2A) with DichroWeb^18,19^ suggesting an *α*-helical fraction of ∼90% for all variants (Table 1). A coiled-coil has a CD_222nm_/CD_209nm_ close to or above 1,^20^ whereas an isolated helix has a stronger negative peak at 209nm. All variants have ratios close to 1 consistent with coiled-coil structures. Furthermore, a SR-CD thermal melt ofAPH_10_ (Fig. S1) showed a clear sigmoidal unfolding curve, suggestion that the coiled-coil unfolded cooperatively.

**Table 1.**
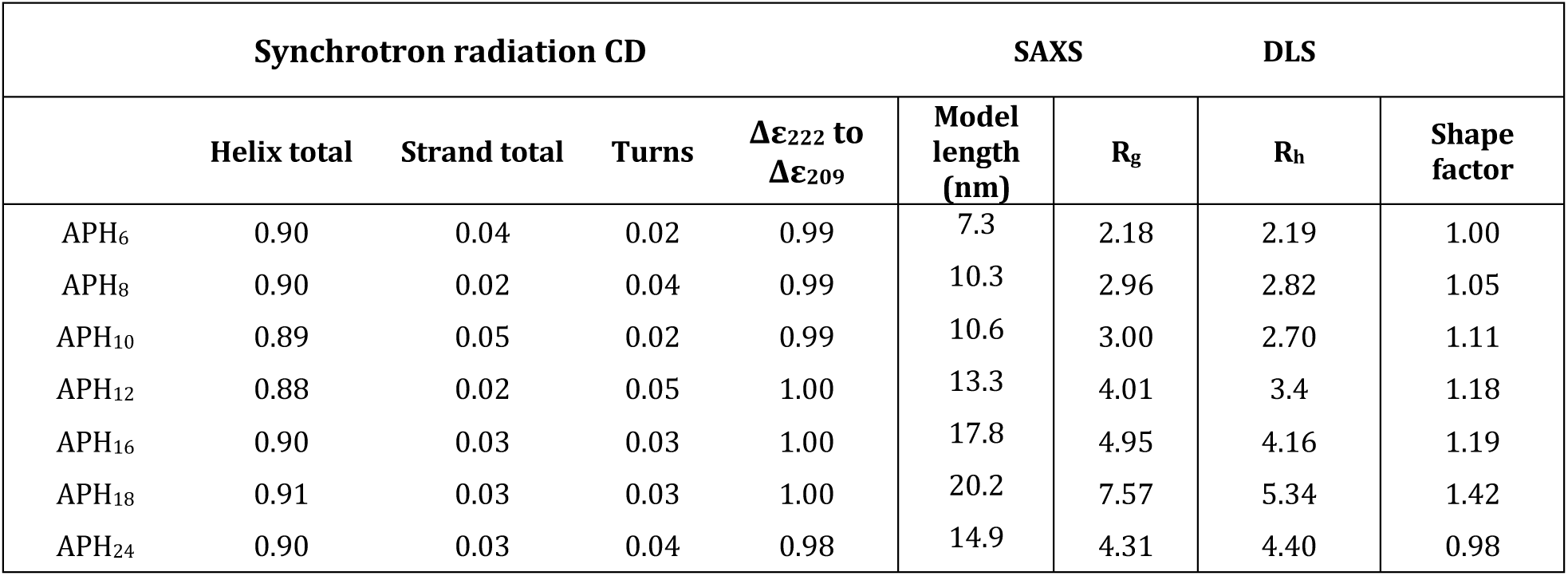
Biophysical parameters for APH series

**Figure 2.**
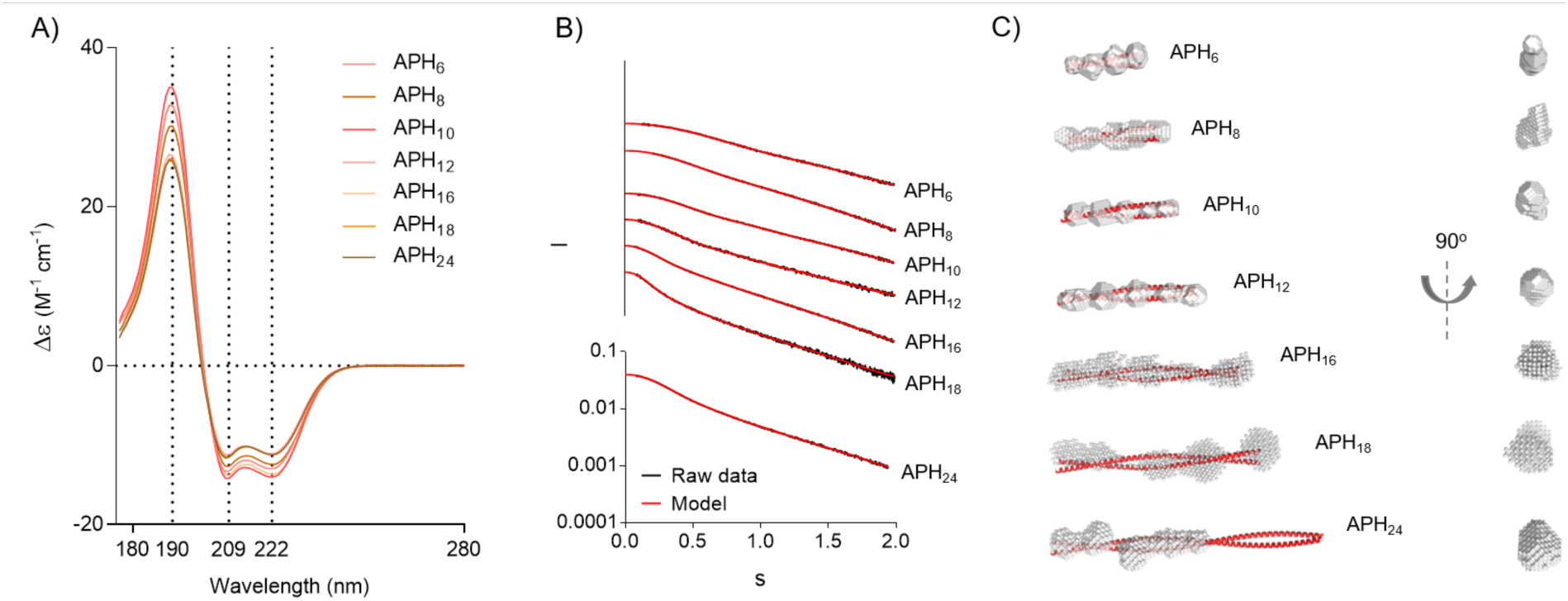
Biophysical characterization of APH variants. Structure of each of the APH spacers was characterized using SR-CD (A) and SAXS (B and C). (A) shows the SR-CD spectra of spacers with four to twenty four heptad repeats measured at 25^*°*^C. Raw SAXS data (shown in black) and data back predicted from the *ab initio* models (shown in red) is summarized in (B). *Ab initio* SAXS models of all APH variants are shown in white with the CCBuilder2.0 models shown as red helixes (C).

To characterize the shape and size of APH variants, they were examined by dynamic light scattering (DLS). All samples could be adequately described by a single diffusion time, which suggested that the samples were monodisperse. As expected, the hydrodynamic radius (R_h_) increased with the number of heptad repeats for all constructs except APH_24_. The R_g_ from SAXS and R_h_ from DLS can be combined into a shape factor. A sphere is expected to have shape factor of 0.77, whereas it increases with length for elongated or unfolded molecules.^24^ For all APH variants except APH_24_, the shape factor increased with the number of heptad repeats, consistent with more elongated structures (Table 1).

The overall shape of the APH variants was studied by small angle X-ray scattering (SAXS) (Fig. 2B, S2-3). The scattering curves were used for *ab initio* model building resulting in elongated shapes for all constructs (Fig. 2C). For coiled-coils containing between 6 and 18 repeats, the lengths of the *ab initio* shape (Fig. 2C, Table 1) matched idealized coiled-coils constructed using CCBuilder2.0.^21^ The cross-section of the *ab initio* shape-models is roughly uniform through-out the series. This suggests these variants form rigid and stable coiled-coils as desired. The longest variant, APH_24_, formed a much shorter structure than the intended design. This could suggest an intra-molecular hairpin, but the molecular weight suggests a dimeric form. APH_24_ does not have the intended structure and will not be discussed further, but will still be included for completeness.

For a more stringent comparison of the SAXS data to the models, we back-predicted the scattering from the idealized coiled-coil models using CRYSOL,^22^ and compared them to the experimental scattering (Fig. S4). APH6 and APH12 show excellent agreement with the back-predicted scattering curve, suggesting that the nanocalipers that their structure is fully described by the idealized model. For APH_8_, APH_10_, APH_16_ and APH_18_ there are small, but notable deviations from the back-predicted curve. This could be due to bending, unfolding, or a minor population of larger complexes. The molecular weight calculated from SAXS data match expectations for a homodimer for APH_6_, APH_10_ and APH_12_, whereas APH_8_, APH_16_ and APH_18_ have 22-37% higher molecular weights than expected. Molecular weights determination by SAXS is less reliable for highly extended molecules,^23^ but the molecular weight deviation may indicate larger complexes forming at high protein concentrations. This is especially pronounced for the constructs that are destabilized by removal of repeats from the consensus design such as APH_8,_ or are very long such as APH_16_ or APH_18_. Notably, the binding experiments are done at protein concentrations that are at least 100-fold lower, and do not show any sign of the large avidity effects expected for stable higher order complexes. Thus, it is unlikely that such transient higher order structure have a major impact on the binding experiments. APH_10_ has the expected molecular weight, but show a small deviation from the predicted scattering for an idealized coiled-coil, however, the *ab initio* modelling suggested that these structures have the intended average dimensions. This leaves bending of the coiled-coil as the most likely explanation, which is unlikely to have a major influence on the inter-epitope distance. In total, data from SR-CD, SAXS and DLS showed that the APH-variants with between 6 and 18 repeats mainly form the desired rigid, coiled-coil domains.

To estimate the distance between the epitopes, we modelled the flexible His-tags on the idealized coiled-coil nanocalipers using EOM.^25^ The distribution of distances was measured between the third histidine of each tag (Fig. S5). For each nanocaliper, the inter-epitope forms a narrow distribution with a mean about a nm longer than the coiled-coil. The successful designs of APH_6_ to APH_18_ thus span a mean inter-epitope distance from 8 to 21 nm, which match the expected intra-Fab distances in an antibody. This suggest that these proteins can be used to probe the spatial dependence of avidity by acting as nanocalipers.

As the avidity effect is mainly on the dissociation rate, we used the surface plasmon resonance technique to study the interaction as it allows accurate quantification of tight interactions that vary across orders of magnitude in stability. Furthermore, low-density immobilization of the antibody on a surface prevents formation of higher order complexes. Such higher order complexes are likely to confound analysis of solution experiments. Initial screening of several commercial anti-His-tag antibodies suggested that THE His-antibody from Genscript performed best in terms of affinity and monodispersity. This antibody is a murine IgG1 antibody with native hinge sequences, and can thus be used as a model system for its class. The oriented capture system used here leaves both binding sites exposed for the interaction with antigens (Fig. 1D). Bivalent binding occurs in two steps: An initially monovalent association is followed by an intra-complex ring-closing reaction to form the cyclical complex (Fig. 1B). As SPR measures change in the refractive index corresponding to the mass accumulated on the chip surface, the ring-closing reactions cannot be directly observed. However, dissociation from the antibody require simultaneous dissociation of both sites. When avidity occurs, dissociation is thus slower from bivalent complexes, and can be quantified as the enhancement factor β defined by the ratio between k_off_mono_ with bivalent k_off_.^26^

To quantify avidity, we measured the interaction kinetics of a monoclonal antibody with mono- and bivalent APH variants. SPR data recorded for APH_half_ could be fitted to a 1:1 interaction model (Fig 3A-C). As expected, the affinity is pH sensitive and changes from a K_D_ of 340 ± 9 nM at pH 5.8 to 40 ± 0.2 nM at pH 6.2 (Table S1). The pH range was not extended further, because lower pH lead to rapid deterioration of the Fc capture system, whereas high pH lead to unpractically slow dissociation of bivalent complexes. The dissociation of APH_12_ was biphasic at all pH values (Fig. 3D-F). The rapid component had a dissociation rate similar to that observed for monovalent complexes. This suggested that the fast phase represented proteins bound only to a single site, while the slower phase corresponded to dissociation of bivalently bound protein. The biphasic behavior was more pronounced at higher concentrations, likely because the binding of a second APH dimer to one antibody is concentration-dependent, and thus compete more efficiently with the intra-molecular ring-closing reaction at high concentrations (Fig. S6A). The kinetics could be described well by a global two-component binding model, which resulted in a fast k_off1_ similar to k_off_mono_ and a slower k_off2_. In all case, the fitted second association rate, k_on2_, was vanishingly small (Table S1), which suggested that the system was well-described by a single association. The two-component model used in the BIACORE Evaluation software did not fully capture the concentration dependent amplitudes and produced unrealistically small error estimates. Therefore, we refitted the dissociation rate separately with global rate-constants and variable amplitudes, and used this to estimate the confidence intervals from the *χ*^2^-surface (Fig. S7). Both analyses produced similar values (Fig. S7A), but the separate fit of the dissociation phase resulted in a slightly better fit, and these dissociation rates will be discussed in the following.

**Figure 3.**
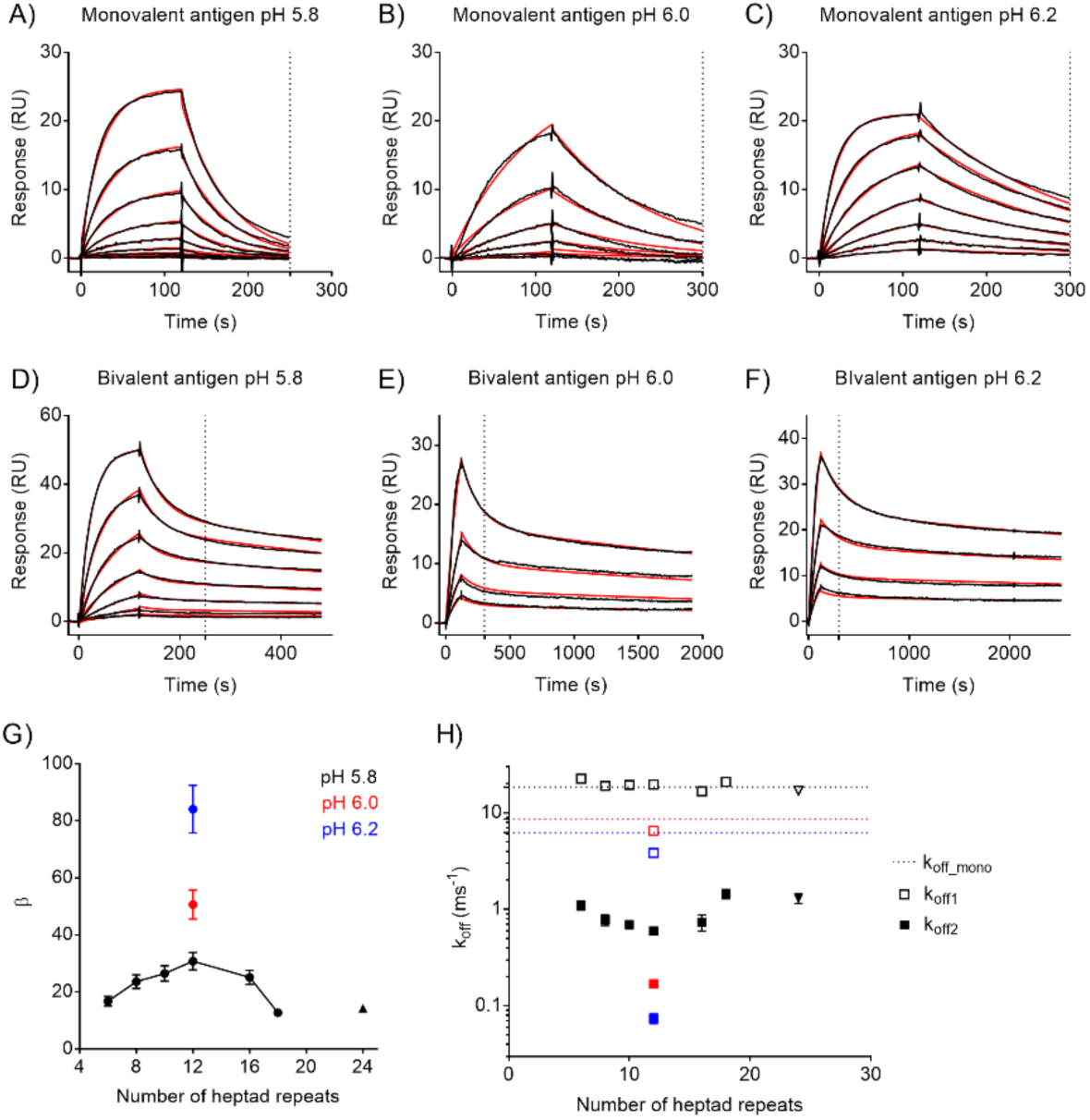
Evaluation of avidity enhancement by surface plasmon resonance. Representative sensorgrams of APH_half_ (monovalent) and APH_12_ (bivalent) binding to the anti-His antibody at various pH are shown in (A) to (F). 250 sec (A and D) or 300 sec (B, C, E and F) are marked. Each line represents a single antigen concentration from the dilution series injected over the chip surface. Raw data is shown in black with 1:1 (A, B and C) or bivalent analyte (D, E and F) fit shown in red. The bivalent binding produces a biphasic dissociation curve with a fast component (k_off1_) coming from the monovalent interaction and slow component (k_off2_) coming from the bivalent interaction (k_off_bi_). The affinity enhancement (β) expressed as a fold decrease in the k_off_bi_ as compared with k_off_mono_ determined for monovalent antigen is shown in (G). The log(k_off_) values of all antigen variants are summarized in (H) with log(k_off1_) and log(k_off2_) are shown as open and full symbols respectively. k_off_mono_ is the dissociation rate constant of the monovalent antigen (APH_half_).

To map the spatial dependence of avidity, we measured binding kinetics for all bivalent APH variants (Fig. S5). The complete series was measured at pH 5.8, as the K_D_ was sufficiently low to allow precise determination of k_off2_ with relatively short dissociation time. All variants showed dissociation traces similar to APH_12_ (Fig. S6). We recorded the kinetics for APH_24_, but will not interpret these since they do not form the desired structure. There is no apparent trend in the variation in association rates. The association phase is affected by the silent ring-closing step that is not modelled in the fitting, and we thus do not believe that the association rates are well-constrained by the data. In contrast, the dissociation rates shows a smooth dependence on nanocaliper length. In the following, we analyze the avidity enhancement β from complex stability only as defined previously (β = k_off_mono_/k_off2_).^27^ Comparison of the k_off2_ to k_off_mono_ for all nanocalipers allowed us to determine the avidity enhancement at epitope spacings ranging from 8 to 21 nm (Fig. 3G). The avidity enhancement reached a maximum at a spacing of 14 nm (30-fold). An increase of the spacing to 21 nm resulted in a two-fold decrease to a β of 12.8, where as a decrease to 8 nm lead to a decrease to a β of 16.8. This suggested that the spatial variation of avidity was relatively small. While β-values are intuitively understandable, avidity is best understood in terms of free energy (Fig. 3H). The two-fold difference in k_off_ suggests that structural perturbations occurring in the antibody ensemble upon stretching are only on the order of 400 cal/mol. This shows that the energy landscape for avidity is relatively flat for IgG1 antibodies. This suggests that the antibody scaffold has been selected to provide avidity in multivalent antigens with a wide range of spatial arrangements. In total, our results describes a flat energy landscape where the exact distance between antigens has a relatively little influence within the spatial tolerance of the antibody.

During the preparation of this manuscript, two elegant studies appeared that also aimed at describing the spatial dependence of avidity. Using DNA origami to control the spacing of epitopes,^28^ Shaw *et al*. found that maximal avidity occurred at a spacing of 16 nm, and decreased ten-fold at a spacing of 17 nm. This study also suggested that the spatial dependence of avidity depended strongly on the strength of the monovalent affinity.^28^ At a monovalent K_D_ ∼3 μM, bivalent binding and avidity enhancement was only observed at the optimal epitope spacing, whereas avidity was observed at a broad range of spacings at a monovalent K_D_ ∼30 nM. Similarly, Zhang et al. used triangular DNA nanostructures and high-speed AFM to measure multivalent binding to epitopes spaced from 3 to 20 nm.^29^ Optimal avidity was observed at a spacing of 10 nm, but with similar values at spacings of 8 and 16 nm, whereas spacings below 5 and above 20 nm did not give stable bivalent binding. As the goal is to extract a consensus description of avidity in antibodies, it is worthwhile to compare the similarities and differences between these studies.

Our study was designed to take advantage of the pH dependence of the His-tag binding to tune the monovalent K_D_. We have examined the spatial dependence on avidity at pH 5.8, where we see a monovalent K_D_ of 320 nM. In free energy terms, this is in the middle of the two affinities used by Shaw *et al*., while Zhang et al. do not report affinities. We observed avidity at a broad range of epitope spacings in agreement with conclusion reached previously for the higher affinity antigen. In principle, the pH sensitivity of the His-tag allow us to test a wider range of monovalent affinities. In practice, the range of pH values probed is limited by the stability of the capture system at low pH, and impractical long dissociation times at higher pH. The current design is limited to pH values ranging from 5.8-6.2, which still results in a ∼3-fold decrease in the monovalent k_off_. Theory suggests that for a bivalent interaction, the combined affinity is given by:^30^

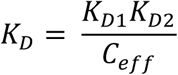

Where K_D1_ and K_D2_ are the monovalent affinities of the two interactions which are identical here, and c_eff_ is the effective concentration of ring-closing. Alternatively, the strength of a bivalent interaction can be decomposed into twice the free energy of the monovalent interaction, and a “coupling free energy” determined by the molecular architecture.^17^ These formulations are equivalent as effective concentration determines the coupling free energy. Both formulations thus predict that the avidity enhancement should increase with the strength of the monovalent interaction. In agreement with this, the avidity enhancement increases from 30-fold to 51- and 84-fold (Fig. 3G) at pH 6.0 and 6.2, respectively. This is roughly consistent with the additivity of the monovalent interaction energies, which predicts that bivalent interaction should have double the pH dependence of the monovalent in energetic terms. The additivity is not perfect, which is frequently observed in biological interactions due to structural compensation.^32^ The assumptions underlying constant “coupling free energy” assumes full ring-closing. It is thus expected that avidity should cease when the effective concentration of the ring-closing falls below the K_D_ of the monovalent interaction,^27^ which explains why the avidity enhancement eventually cease at lower monovalent affinities.

In antibody engineering, it is crucial to determine both the spacing providing the maximal avidity enhancement and the total tolerable spacing for bivalent binding. We observe a maximal avidity at a spacing of 14 nm, which is between the optimal spacings reported by *Zhang* and Shaw *et al*. of 10 and 16 nm respectively. Furthermore, we still see bivalent binding with little destabilization at a spacing of 20nm, whereas Shaw *et al*. observed at sharp drop in stability at 17nm and Zhang does not observe bivalent binding at 21 nm. Ultimately, the distance dependence of avidity is determined by the structure and flexibility of the antibody and is likely to be similar but not identical for different antigens. There are several differences between these studies that could explain the differences including the antibodies/antigens, the epitope-linking chemistry, the immobilization strategy and the nature of the nanocaliper. Of these factors, the flexibility of the epitope relative to the nanocaliper is likely the biggest contributor.

An flexible epitope can extend beyond the length of the nanocaliper, or fold back along the length of the coiled-coil. Similarly, it determines the allowed domain conformations of the bound Fab moiety. Thereby a flexible antigen attachment smoothen the distance dependence and increase the apparent reach of the antibody. The three-residue linker used here only suffice for a tight turn and thus likely represent a minimal flexible connection. An extended six-residue His-tag is about 2 nm long, and back-folding of the His-tag towards the center of the coiled-coil would shorten the distance. In practice, this shortening is likely less than the theoretical maximum due to the strain imposed by binding orthogonal to the coiled-coil. At the longest extensions, it is likely that the antibody binds in a T-shaped conformation with the Fab moiety parallel to the coiled-coil. In such a conformation, the flexibility of the linker used here is unlikely to contribute significantly to the epitope distance. This suggest that the coiled-coil system developed here likely provide a good estimate of the energetics of antigen spacing near the maximal extension of the antibody. At small epitope spacings, it is likely that the His-tag can be bound in a conformation where they are stretched out from the end of the coiled-coil. Therefore, the size of the epitope itself contributes significantly to the spacer, and it may be inherently difficult to probe the minimal acceptable spacing with flexible epitopes. Therefore, we did not pursue shorter coiled-coil constructs further.

The epitopes used in this study are flexible relative to the nanocaliper, and thus have comparable rotational freedom as the small molecules flexibly conjugated to a surface used previously.^28,29^ The rotational freedom of the epitope removes the potential contribution from inter-epitope orientation, which is likely to restrict the range the antibody spans as the Fab fragments have to bind in certain orientations. The distance dependence of fixed-angle epitopes is thus likely not well-modeled by the system developed here. However, many multivalent drug targets allow epitopes to rotate relative to each other, for example membrane proteins diffusing in a membrane or protein complexes with multiple domains or subunits. Freely rotating epitopes thus represent a sizeable fraction of antibody targets that are likely to have a similar distance dependence.

The spatial dependence of avidity described here is qualitatively different from that described by Shaw *et al*.. In terms of an energy landscape, Shaw *et al*. suggest maximal avidity is achieved at the foot of a steep slope, where a slight increase of the distance is very detrimental to binding. We suggest that the optimum avidity is achieved at the bottom of a broad valley, where the precise spacing is not critical as long as it is within the spatial tolerance of the antibody. For the rational use of avidity in antibody engineering, this is a critical difference. The latter view was also supported by a recent study which suggested a relatively weak avidity dependence at epitope spacings between 8 and 16 nm.^29^ The energy landscape of avidity is determined by a combination of inter- and intramolecular interactions. Within the spatial range where bivalent interaction can occur without distorting the binding sites, the avidity landscape is likely to be dominated by the structural ensemble of the antibody. Structural studies of isolated antibodies suggest that they sample a wide variety of conformations with no particular preferred interdomain orientations. This suggests that most interdomain orientation in antibodies are of similar energy, which is consistent with a wide and flat energy landscape of avidity. For such a flexible bivalent interaction, statistical thermodynamics suggests that optimal avidity should occur when the spacing of the epitopes match the average spacing between binding sites in the flexible antibody.^10^ The average spacing is necessarily far from the maximal extension of the antibody. Maximal avidity is thus only likely to occur near the maximal extension of the antibody if new inter- or intra-molecular interactions are formed in the bound state. Such interactions have been observed previously and are likely to be system specific.^5^ We thus believe the case observed here, where maximal avidity is found at the center of a broad valley represent the more generic case.

The protein nanocaliper system based here has advantages and disadvantages compared to previous systems based on DNA origami. Peptide antigens can readily be encoded into a protein-based system developed here, which is crucial as many antigens are proteins. Furthermore, the spatial resolution of controlled antigen presentation is inherently determined by the pitch of the helix. A coiled-coil based system thus in principle allow controlled spacings down to a single turn of a *α*-helix (∼5Å), where the local environment surrounding the epitope is identical. The larger turn of a DNA double helix (34Å) may cause additional contributions from the orientation of the epitope when close epitope spacings are closed. Furthermore, while we do experiments on a surface attached system, the coiled-coil system developed is immediately compatible with solution assays. The main drawback of a coiled-coil based nanocaliper system is the refolding efficiency of the coiled-coil structure decrease strongly with length, likely because the repeating coiled-coil fold is prone to kinetic trapping. Although it would have been desirable to reach a length where bivalent binding is abrogated, the simple design approach we have used here, can likely not be extended much further. In contrast, DNA origami can be used to create much larger structures, and much more complex geometric patterns. Both DNA- and protein-based nanocaliper systems have advantages, and we believe that the system developed here represent a useful complement to existing DNA-based technologies.

Avidity holds great promise to enhance the affinity and specificity of target recognition by antibodies. Bivalent binding is particularly important in bispecific antibodies, where Fab moieties that bind different epitopes are combined. In such systems, simultaneous engagement of the epitopes is crucial to the desired therapeutic effect. Likely, nanocaliper based control of the epitopes spacing will play a key role in elucidating the spatial and energetic landscape that underlies bivalent binding, and may help develop new antibody formats that optimize avidity.

## Acknowledgements

We would like to thank Cy M. Jeffries for the assistance in using the Petra III beamline, Nykola Jones for assistance in SR-CD experiments, and Michael Ploug and Gregers R. Andersen for critical comments to the manuscript. This work was funded by a grants from the Villum Young Investigator program, The Independent Research Fund Denmark and the Marie Curie COFUND program. Access to the SR-CD beam line was granted by ISA, Centre for Storage Ring Facilities, Aarhus at Aarhus University.

## Materials and methods

### Preparation of DNA constructs

Synthetic DNA of APH constructs were purchased from Gene Universal (Newark, USA) and GenScript (Piscataway, USA) cloned into a pET15b expression vector. The protein variants contained a 6xHis-tag followed by a short flexible liker (GSS) before the N-terminus of the coiled-coil. A thrombin site was introduced between the 6xHis-tag and the coiled-coil using the QuikChange Lightning mutagenesis kit (Agilent, Santa Clara, USA). Sequences of the products were confirmed by DNA sequencing. Sequences of all variants are given in the supplementary material.

### Expression, purification and refolding of APH variants

All APH variants were expressed in inclusion bodies in *E. coli* BL21 (DE3). Cells were cultured over-night at 37^°^C in autoinduction medium^33^ supplied with 100 µg/ml ampicillin and harvested by centrifugation (8983g, 20 min, 4^°^C). Cell pellets were resuspended in lysis buffer containing 50 mM Tris-HCl pH 8.0, 200 mM NaCl, 1 mM EDTA, 1% Triton X-100, incubated on ice for approx. 1 h and disrupted by sonication. Insoluble fraction containing inclusion bodies was then isolated by centrifugation (14000 g, 4^°^C, 30min) and washed in 20 mM Na_2_HPO_4_, 0.5 M NaCl, pH 7.4 by gentle stirring for 30 min at 4°C followed by removal of the buffer by centrifugation (14000 g, 30min, 4^°^C). Inclusion bodies were solubilized in 20 mM Na_2_HPO_4_, 0.5 M NaCl, 6 M urea pH 7.4 by over-night gentle stirring at 4^°^C followed by centrifugation (14000 g, 30min, 4^°^C) to remove insoluble cell debris. APH variants were purified from urea-resuspended inclusion bodies in the presence of 6 M urea on a gravity Ni-sepharose (GE Healthcare) column and eluted with the step gradient including imidazole concentrations 5 mM, 20 mM, 40 mM, 60 mM, 80 mM, 100 mM and 500 mM.

Fractions containing antigens were diluted in buffer containing 6 M urea to a protein concentration of 0.5-1 mg/ml and refolded by stepwise removal of a denaturation agent. For APH_6_, APH_8_, APH_10_ and APH_12_, refolding included three dialysis steps in 20 mM Na_2_HPO_4_, 66.7 mM NaCl, pH 7.4 at 4^°^C: I) 3 M urea, overnight. II) 1 M urea, ∼8h. III) no urea, overnight. For APH_16_, APH_18_, and APH_24_, refolding included four dialysis steps I) 4.5M urea, overnight. II) 2.5 M urea, ∼ 8h. III) 1 M urea, over-night. IV) no urea, ∼8h. Misfolded proteins were removed by centrifugation (14000 g, 4^°^C, 30min), heating to 80^°^C for 20 min, followed by centrifugation to remove proteins that were not thermostable. Supernatants were purified on the Superdex75 10/300 GL column (GE Healthcare) (APH_6_, APH_8_, APH_10_, APH_12_) or Superdex200 10/300 GL (GE Healthcare) (APH_16_, APH_18_, and APH_24_) equilibrated in 20 mM Na_2_HPO_4_, 0.3 M NaCl, pH 7.4. Size exclusion of APH_24_ was additionally followed by ion exchange chromatography on Source S15 4.6/100 PE (GE Healthcare) column in buffer containing 20 mM Na_2_HPO_4_ pH 7.4. APH_24_ was eluted with a linear gradient of 0-1M NaCl over 10 CV. These purification procedures resulted in a protein of more than 95% purity as judged by SDS-PAGE.

The samples used for SAXS, SR-CD and DLS contained a thrombin-site to allow removal of the His-tag. These proteins were purified identically to the variants without the cleavage site. The purified protein was incubated and for 1h at 37^°^C with bovine thrombin (Sigma) in a 1000:1 molar ratio. Thrombin was then removed by heating to 80^°^C for 20 min followed by centrifugation.

### Synchrotron radiation circular dichroism (SR-CD)

All spacers with removed His-tag were analyzed by SR-CD at AU-CD beam line in ASTRID2 storage ring, Aarhus University, Aarhus, Denmark. In order to remove Cl^-^ ions all spacers were dialyzed against 20 mM Na_2_HPO_4_, 0.15 M NaF, pH 7.4 prior SR-CD measurements. Concertation of each spacer was brought to approx. 0.7-1 mg/ml what resulted in absorbance below 1. APH spacers were scanned three times over 280 to 170 nm at 25^°^C using quartz cuvette with 0.102 mm pathlength.^34^ Reference spectrum from the buffer was obtained analogously and was subtracted from the protein spectrum prior data analysis. For visualization all three protein spectra were averaged and mildly smoothened with a 7 point Savitzky-Golay filter.^35^ To compare different linker lengths spectra were scaled according to the protein concertation calculated from the absorption at 205 nm. Secondary structure content was calculated using DichroWeb server.^18,19^

Melting experiment was performed for APH_10_. Sample preparation and parameters used for measurement were analogical as for analysis at 25^°^C. Melting was followed at temperatures from 25^°^C to 85^°^C with 5^°^C steps. After reaching each step system equilibrated for 5 min after which protein spectrum was measured over 280 nm to 170 nm. After reaching 85^°^C samples were cooled down to 25^°^C and protein spectrum was measured again.

### Small angle X-ray scattering (SAXS)

The synchrotron SAXS data was collected at beamline P12 operated by EMBL Hamburg at the PETRA III storage ring (DESY, Hamburg, Germany).^36^ All measurements were performed at 20°C in 20 mM Na_2_HPO_4_, 150 mM NaCl pH 7.4. To minimize radiation damage 5 mM DTT was added to each sample and the reference buffer immediately before measurements. Signal from each spacer was obtained for at least three protein concentrations varying from 14 mg/ml to 0.5 mg/ml. As no indication of the concentration dependent scattering was observed model reconstruction was performed for the highest concentration only where signal to noise ratio was the lowest. Scattering from the buffer was subtracted from the raw data and the scattering curves were brought to absolute scale using known scattering cross section of water. Prior analysis data was averaged and normalized to the intensity of the incident beam. The *ab initio* models were build using DAMMIF software,^37^ a part of the ATSAS 2.8.4. package,^22^ using 10 repetitions and P1 symmetry. Models were averaged with DAMAVER^38^ and refined with DAMMIN.^39^ Idealized coiled-coil models were built using CCBuilder2.0,^21^ and the His-tags were modelled as pseudo-atoms in a random coil conformation using EOM.^25^

### Dynamic light scattering (DLS)

All experiments were performed on a Wyatt DynaPro NanoStar DLS instrument at 20^°^C on samples from size exclusion chromatography. The data was analysed using the DYNAMICS software to extract the hydrodynamic radius (R_h_).

### Surface plasmon resonance (SPR)

Affinity of an interaction between the antigen and antibody were determined using SPR. All measurements were performed on Biacore T200 (GE Healthcare) instrument at 25^°^C, flow rate 30 µl/min in 20 mM Na_2_HPO_4_, 0.5 M NaCl, 0.05% Tween20, 0.1% BSA at pH 6.2, 6.0 or 5.8. The high NaCl concentration was necessary to prevent non-specific electrostatic sticking.

The SPR chip was prepared by amine coupling of a polyclonal rabbit anti-mouse IgG (GE Healthcare) on both flow cells of CMD500 chip (Xantec). Immobilization was conducted according to the protocol described by manufacturer and resulted in capture of approx. 8000 RU on both active and reference flow cell. Polyclonal anti-mouse IgG targets Fc region of mouse IgG enabling oriented capture of mouse IgG with both Fab moieties exposed for the antigen binding. To determine the binding affinities between antigens and antibody mouse monoclonal anti-His-tag IgG (clone 6G2A9, The^™^ His tag Ab, GenScript) was captured on the surface of active flow cell to the level of 100-200 RU. All antigens with various spacer length were tested for binding at pH 5.8. In each cycle single concentration of an antigen from 2-fold dilution series (200-1.6nM for bivalent and 800-3.2nM for monovalent antigen) was injected over both flow cells for 120 sec allowing association. Dissociation was then measured for 360 sec of constant buffer injection. Between the cycles chip surface was fully regenerated by three 15 sec injections of low pH buffer containing 10mM glycine pH 1.7.

Non-specific binding was removed from the raw data by subtraction of signal from the parallel experiment performed in the reference flow cell without anti-His-tag antibody. Buffer injection was also subtracted from the raw data (i.e. double buffer referencing). Data was analyzed in Biacore T200 Evaluation Software (GE Healthcare). For antigens forming monovalent interactions (APH_half_ and APH_4_) kinetic constants were determined by fitting 1:1 Langmuir interaction model to the binding curves. For antigens bivalently interacting with an antibody (APH_6_, APH_8_, APH_10_, APH_12_, APH_16_, APH_18_, and APH_24_) bivalent analyte model was fitted to the binding curves using one global set of kinetic parameters. Dissociation of these antigens was biphasic with the fast phase more pronounced at high concentrations. The fast phase likely results from two APH dimers bound monovalently to the same antibody, whereas the slow phase refers to dissociation of bivalent antigen:antibody complex (Figure S2). Representative sensorgrams with the fit are shown in (Figure S2). As the concentration dependence of two phases is not included in any of the Biacore Evaluation models, we fitted the dissociation phase globally to a biexponential decay with shared kinetics rates, but variable amplitudes in IGOR Pro (Wavemetrics). The 95%-confidence interval was estimated from the global fit as χ^2^_min_/χ^2^ > 0.9.^40^ Fitted parameters are summarized in (Table S1).

For high affinity complexes, i.e. APH_12_ at pH 6.0 and pH 6.2, 360 sec were not enough to observe biphasic dissociation and decrease in signal sufficient for k_off_ determination. To precisely determine dissociation rate of those complexes antigen was constantly injected over both flow cells for 120 sec and then buffer was constantly injected for 30 min or 40 min. Representative sensorgrams with the fit are shown in (Figure 3).

