## Supplemental Information for "Nanoscale spatial dependence of avidity in an IgG1 antibody"

### Sequences of APH constructs

**Color code:**

**His-tag** Thrombin site *Flexible linker*

**$APH_{half}$**

MGSS**HHHHHH**SSGMKQLEKELKQLEKELQAIEKQLAQLQKKAQARKKKLAQLKKKLQA*PGSGSGP*MKQLEKELKQLEKELQAIEKQLAQLQWKAQARKKKLAQLKKKLQA

**$APH_6$**

MGSS**HHHHHH**SSGMKQLEKELKQLEKELQAIEKQLAQLQWKAQARKKKLAQLKKKLQA

**$APH_{6\_thrombin}$**

MGSS**HHHHHH**SSGLVPRGSHMKQLEKELKQLEKELQAIEKQLAQLQWKAQARKKKLAQLKKKLQA

**$APH_8$**

MGSS**HHHHHH**SSGMKQLEKELKQLEKELQAIEKQLAQLQKKLQAIEKQLAQLQWKAQARKKKLAQLKKKLQA

**$APH_{8\_thrombin}$**

MGSS**HHHHHH**SSGLVPRGSHMKQLEKELKQLEKELQAIEKQLAQLQKKLQAIEKQLAQLQWKAQARKKKLAQLKKKLQA

**$APH_{10}$**

MGSS**HHHHHH**SSGMKQLEKELKQLEKELQAIEKQLAQLQKKAQARKKKLKQLEKELQAIEKQLAQLQWKAQARKKKLAQLKKKLQA

**$APH_{10\_thrombin}$**

MGSS**HHHHHH**SSGLVPRGSHMKQLEKELKQLEKELQAIEKQLAQLQKKAQARKKKLKQLEKELQAIEKQLAQLQWKAQARKKKLAQLKKKLQA

**$APH_{12}$**

MGSSHHHHHHSSGMKQLEKELKQLEKELQAIEKQLAQLQKKAQARKKKLAQLKKKLQALEKELKQLEKE
LQAIEKQLAQLQWKAQARKKKLAQLKKKLQA

**APH$_{12_thrombin}$**

MGSSHHHHHHSSGLVPRGSHMKQLEKELKQLEKELQAIEKQLAQLQKKAQARKKKLAQLKKKLQALEKE
LKQLEKELQAIEKQLAQLQWKAQARKKKLAQLKKKLQA

**APH$_{16}$**

MGSSHHHHHHSSGMKQLEKELKQLEKELQAIEKQLAQLQKKAQARKKKLQALEKELKQLEKELQAIEKQL
AQLQKKAQARKKKLAQLKKKLKQLEKELQAIEKQLAQLQWKAQARKKKLAQLKKKLQA

**APH$_{16_thrombin}$**

MGSSHHHHHHSSGLVPRGSHMKQLEKELKQLEKELQAIEKQLAQLQKKAQARKKKLQALEKELKQLEKEL
QAIEKQLAQLQKKAQARKKKLAQLKKKLKQLEKELQAIEKQLAQLQWKAQARKKKLAQLKKKLQA

**APH$_{18}$**

MGSSHHHHHHSSGMKQLEKELKQLEKELQAIEKQLAQLQKKAQARKKKLAQLKKKLQALEKELKQLEKE
LQAIEKQLAQLQKKAQARKKKLAQLKKKLQALEKELKQLEKELQAIEKQLAQLQWKAQARKKKLAQLKKK
LQA

**APH$_{18_thrombin}$**

MGSSHHHHHHSSGLVPRGSHMKQLEKELKQLEKELQAIEKQLAQLQKKAQARKKKLAQLKKKLQALEKE
LKQLEKELQAIEKQLAQLQKKAQARKKKLAQLKKKLQALEKELKQLEKELQAIEKQLAQLQWKAQARKKK
LAQLKKKLQA

**APH$_{24}$**

MGSSHHHHHHSSGMKQLEKELKQLEKELQAIEKQLAQLQKKAQARKKKLAQLKKKLQALEKELKQLEKE
LQAIEKQLAQLQWKAQARKKKLAQLKKKLQALEKELKQLEKELQAIEKQLAQLQKKAQARKKKLAQLKKK
LQALEKELKQLEKELQAIEKQLAQLQWKAQARKKKLAQLKKKLQA

**APH$_{24_thrombin}$**

MGSSHHHHHHSSGLVPRGSMKQLEKELKQLEKELQAIEKQLAQLQKKAQARKKKLAQLKKKLQALEKELK
QLEKELQAIEKQLAQLQWKAQARKKKLAQLKKKLQALEKELKQLEKELQAIEKQLAQLQKKAQARKKKLA
QLKKKLQALEKELKQLEKELQAIEKQLAQLQWKAQARKKKLAQLKKKLQA

**Table S1.** Kinetic constants determined by SPR for APH variants binding to THE His Ab.

| | | BIACORE evaluation software | | | | IGOR Pro | |
|---|---|---|---|---|---|---|---|
| pH | | **$k_{on1}$** **$10^3 \times (Ms)^{-1}$** | **$k_{off1}$** **$(ms^{-1})$** | **$k_{on2}$** **$10^{-4} \times (Ms)^{-1}$** | **$k_{off2}$** **$(ms^{-1})$** | **$k_{off1}$** **$(ms^{-1})$** | **$k_{off2}$** **$(ms^{-1})$** |
| 5.8 | $APH_6$ | 95.8 ± 0.5 | 18.5 ± 0.2 | 2.74 ± 0.02 | 0.80 ± 0.01 | 22.6 ± 1.3 | 1.1 ± 0.12 |
| | $APH_8$ | 76.7 ± 0.4 | 13.7 ± 0.2 | 1.63 ± 0.02 | 0.63 ± 0.02 | 18.9 ± 1.0 | 0.78 ± 0.10 |
| | $APH_{10}$ | 122.5 ± 0.2 | 14.8 ± 0.1 | 2.39 ± 0.01 | 0.66 ± 0.01 | 19.5 ± 1.1 | 0.70 ± 0.068 |
| | $APH_{12}$ | 100.8 ± 0.4 | 19.9 ± 0.2 | 1.74 ± 0.01 | 0.62 ± 0.01 | 19.6 ± 1.1 | 0.60 ± 0.050 |
| | $APH_{16}$ | 29.0 ± 0.3 | 20.8 ± 0.6 | 0.94 ± 0.02 | 0.77 ± 0.03 | 16.7 ± 1.4 | 0.74 ± 0.14 |
| | $APH_{18}$ | 47.7 ± 0.3 | 19.5 ± 0.3 | 1.16 ± 0.01 | 1.26 ± 0.02 | 20.9 ± 1.7 | 1.5 ± 0.16 |
| | $APH_{24}$ | 65.2 ± 0.4 | 21.4 ± 0.4 | 1.08 ± 0.01 | 1.48 ± 0.02 | 17.0 ± 1.2 | 1.3 ± 0.15 |
| | $APH_{half}$ | 54.6 ± 1.4 | 18.5 ± 0.1 | n.d. | n.d. | n.d. | n.d. |
| 6.0 | $APH_{12}$ | 38.7 ± 0.3 | 4.30 ± 0.13 | 0.54 ± 0.01 | 0.18 ± 0.003 | 6.5 ± 0.6 | 0.17 ± 0.0135 |
| | $APH_{half}$ | 118.8 ± 0.5 | 8.61 ± 0.06 | n.d. | n.d. | n.d. | n.d. |
| 6.2 | $APH_{12}$ | 66.9 ± 0.1 | 3.76 ± 0.02 | 0.72 ± 0.004 | 0.11 ± 0.001 | 3.8 ± 0.3 | 0.074 ± 0.0085 |
| | $APH_{half}$ | 152.7 ± 0.7 | 6.22 ± 0.02 | n.d. | n.d. | n.d. | n.d. |

**Table S2.** Molecular weight of APH variants calculated from the sequence of the monomer ($MW_{sequence\_monomer}$) and determined by SAXS ($MW_{SAXS}$).

| | $MW_{sequence\_monomer}$ (kDa) | $MW_{SAXS}$ (kDa) |
|---|---|---|
| $APH_6$ | 5.7 | 10.0 |
| $APH_8$ | 6.2 | 17.0 |
| $APH_{10}$ | 9.0 | 18.0 |
| $APH_{12}$ | 10.6 | 21.0 |
| $APH_{16}$ | 13.9 | 34.0 |
| $APH_{18}$ | 15.5 | 41.0 |
| $APH_{24}$ | 20.4 | 37.0 |

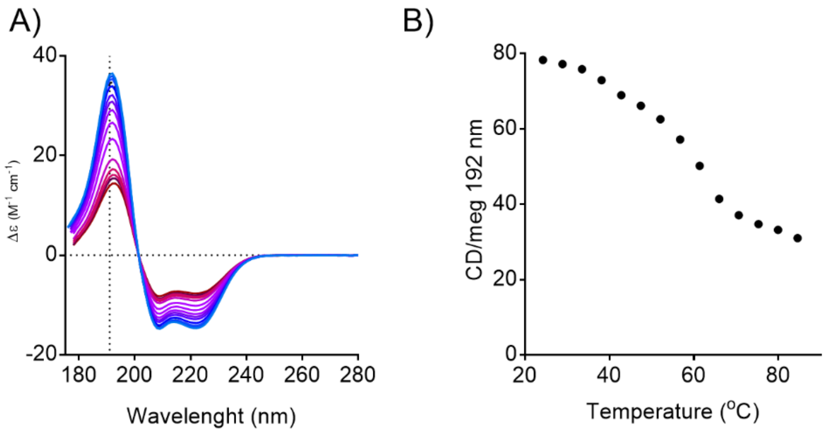


**Figure S1.** SR-CD spectra of $APH_{10}$ were measured at temperatures varying from 25 °C to 85 °C. Dotted line indicates 192 nm the wavelength at which change of the CD signal intensity with temperature was compared.

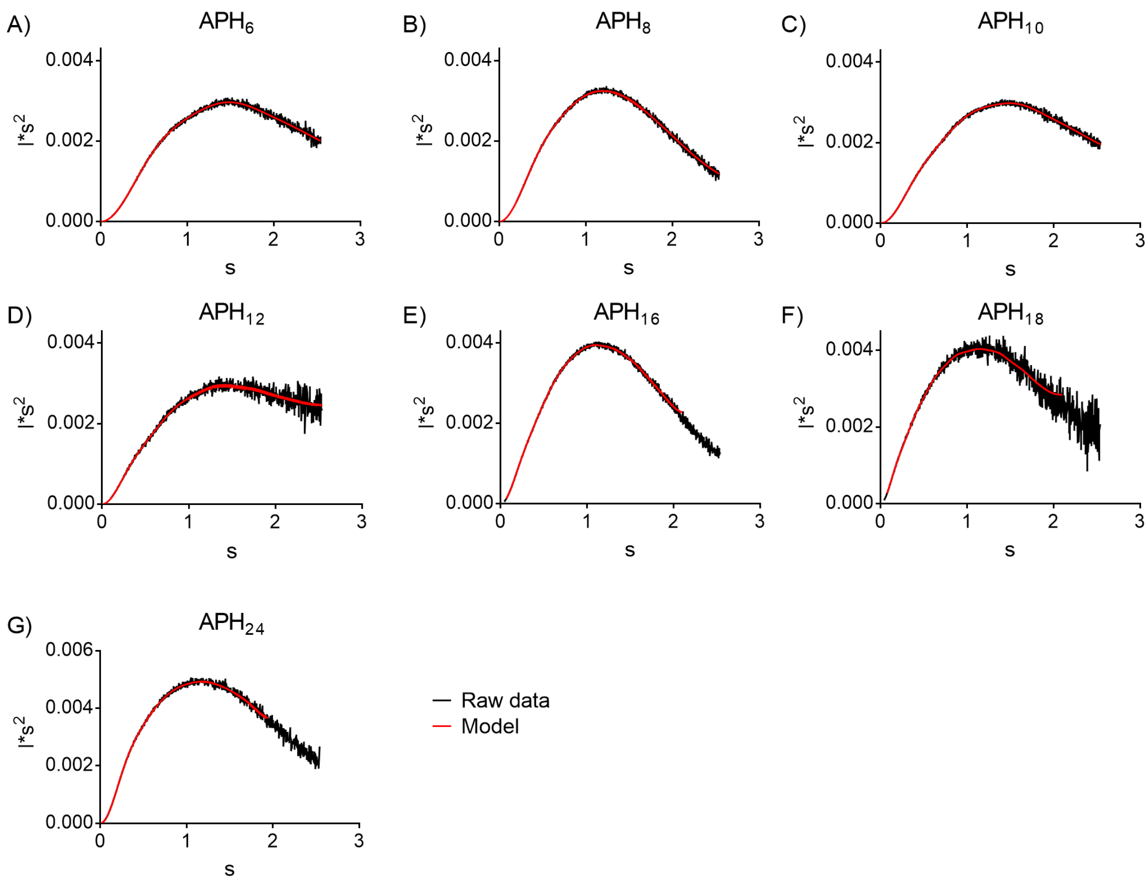


**Figure S2.** SAXS data of all APH variants represented as Kratky plots. Scattering back-predicted from the *ab initio* models represented a red line.

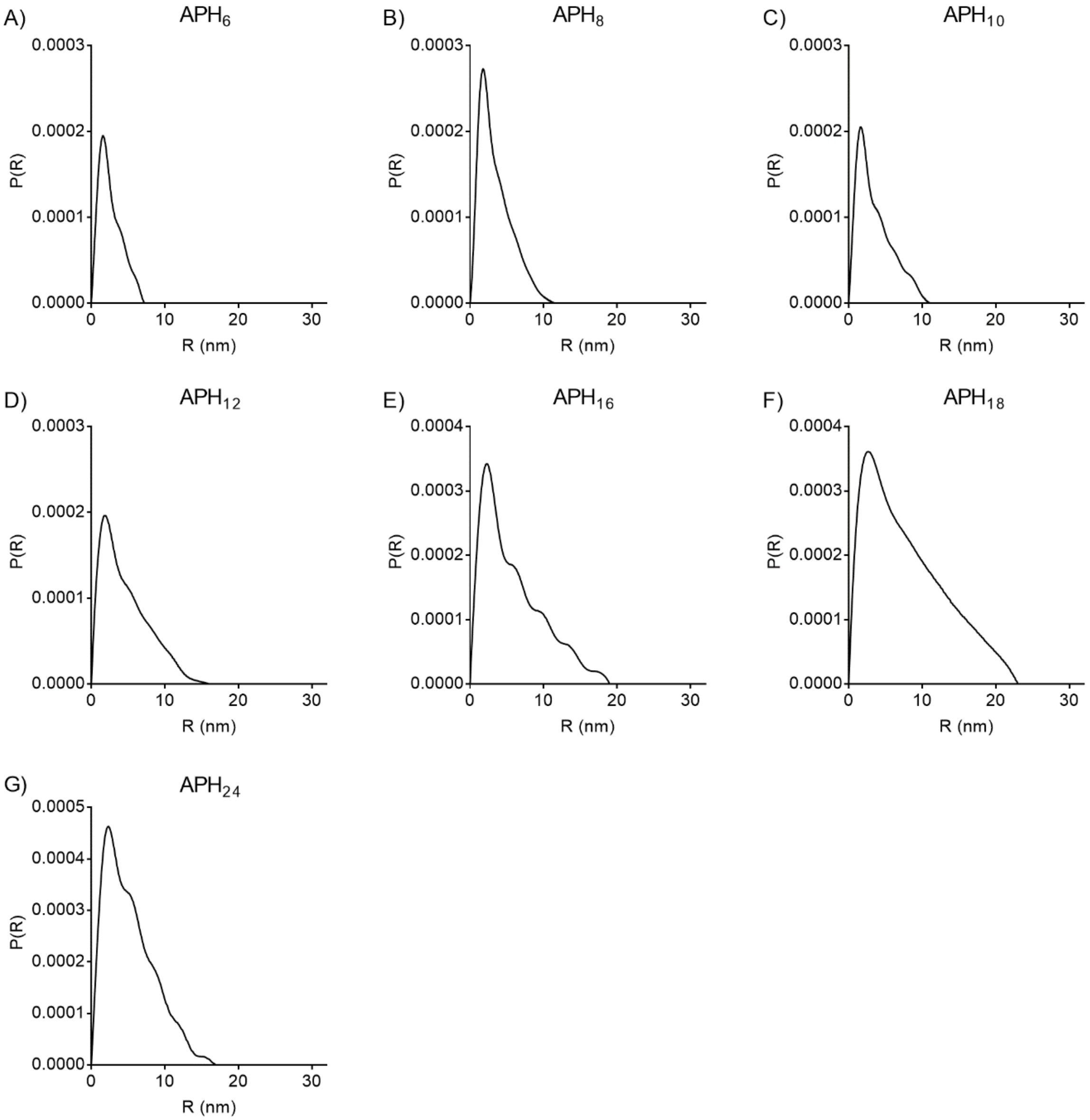


**Figure S3.** SAXS data for all variants represented as p(r) functions. The maximum distance ($D_{max}$) increases steadily up to $APH_{18}$, whereafter it dramatically increase in $APH_{20}$ and is reduced in $APH_{24}$. These data mirror the $R_g$ and the $R_h$ determined from DLS.

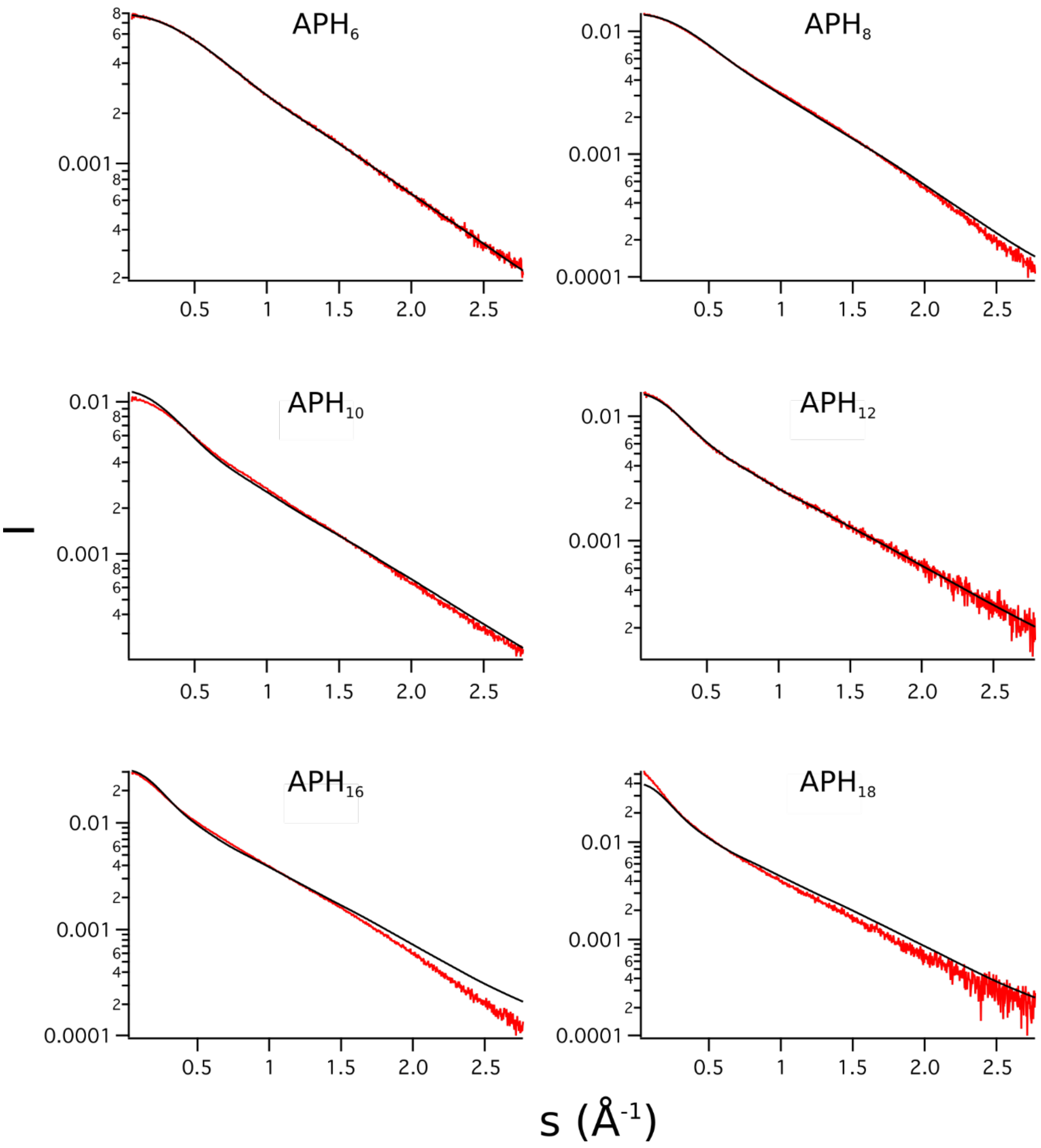


**Figure S4.** Comparison of experimental SAXS measurements (red) to back-predicted scattering curves from idealized coiled-coil models (black).

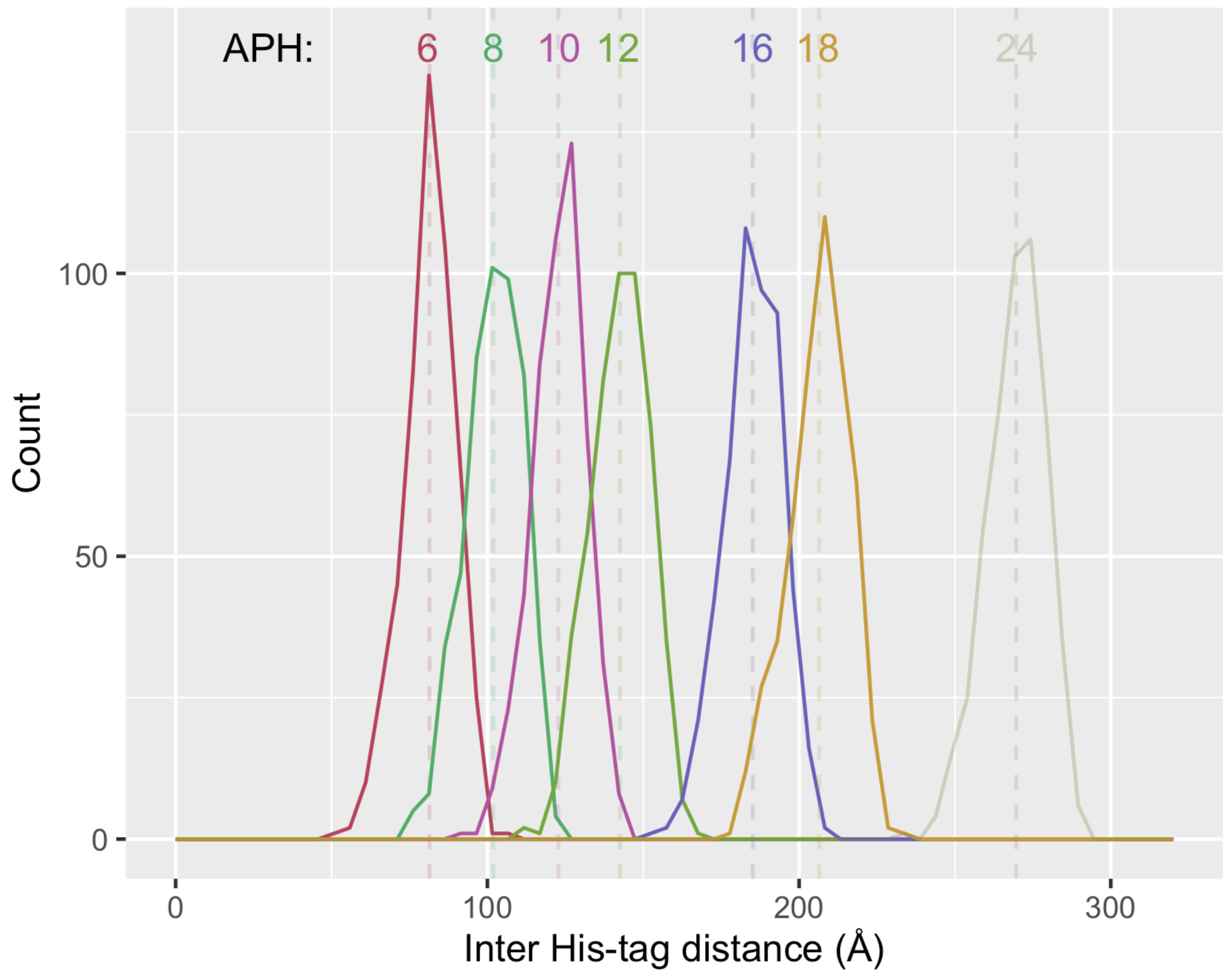


**Fig. S5: Modelled inter-epitope distances in APH nanocalipers. An ensemble representing t**he missing residues were modelled with his-tag in 500 random coil conformations. The inter-epitope distance is measured between the third histidine in each tag. The histogram uses 5 Å bins and the mean distance shown with dashes. Nanocalipers that did not form the desired structure are translucent.

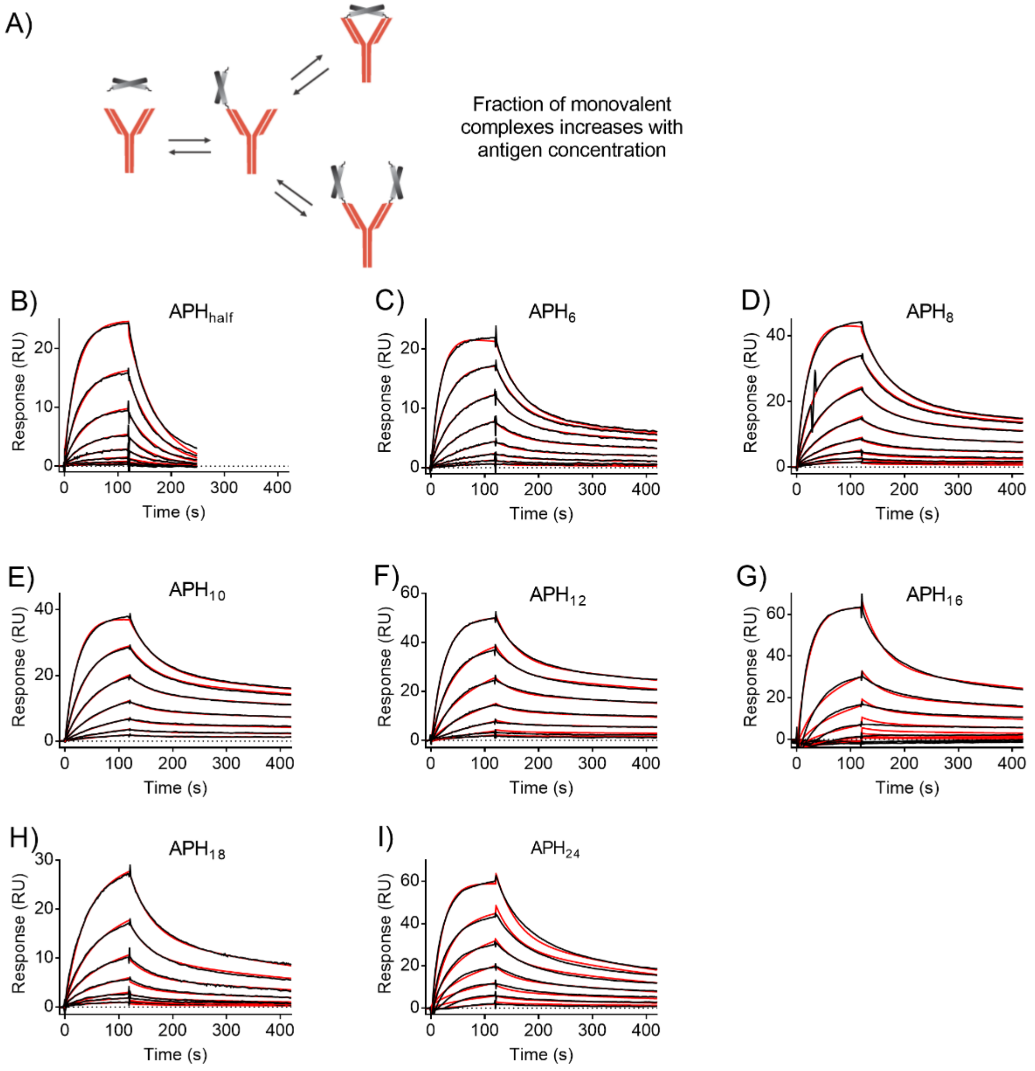


**Figure S6.** Binding of a bivalent antigen to an antibody can result in formation of mono- and bivalent complexes (A). Fraction of antigens interacting with an antibody via single epitope increases with the antigen concertation. Representative sensorgrams of all antigens binding to anti-His antibody at pH 5.8 are shown in (B) to (K). Black lines refer to raw data and 1:1 ($APH_{half}$) or bivalent analyte ($APH_6$, $APH_{10}$, $APH_{12}$, $APH_{16}$, $APH_{18}$, $APH_{24}$) fit is shown as red lines.

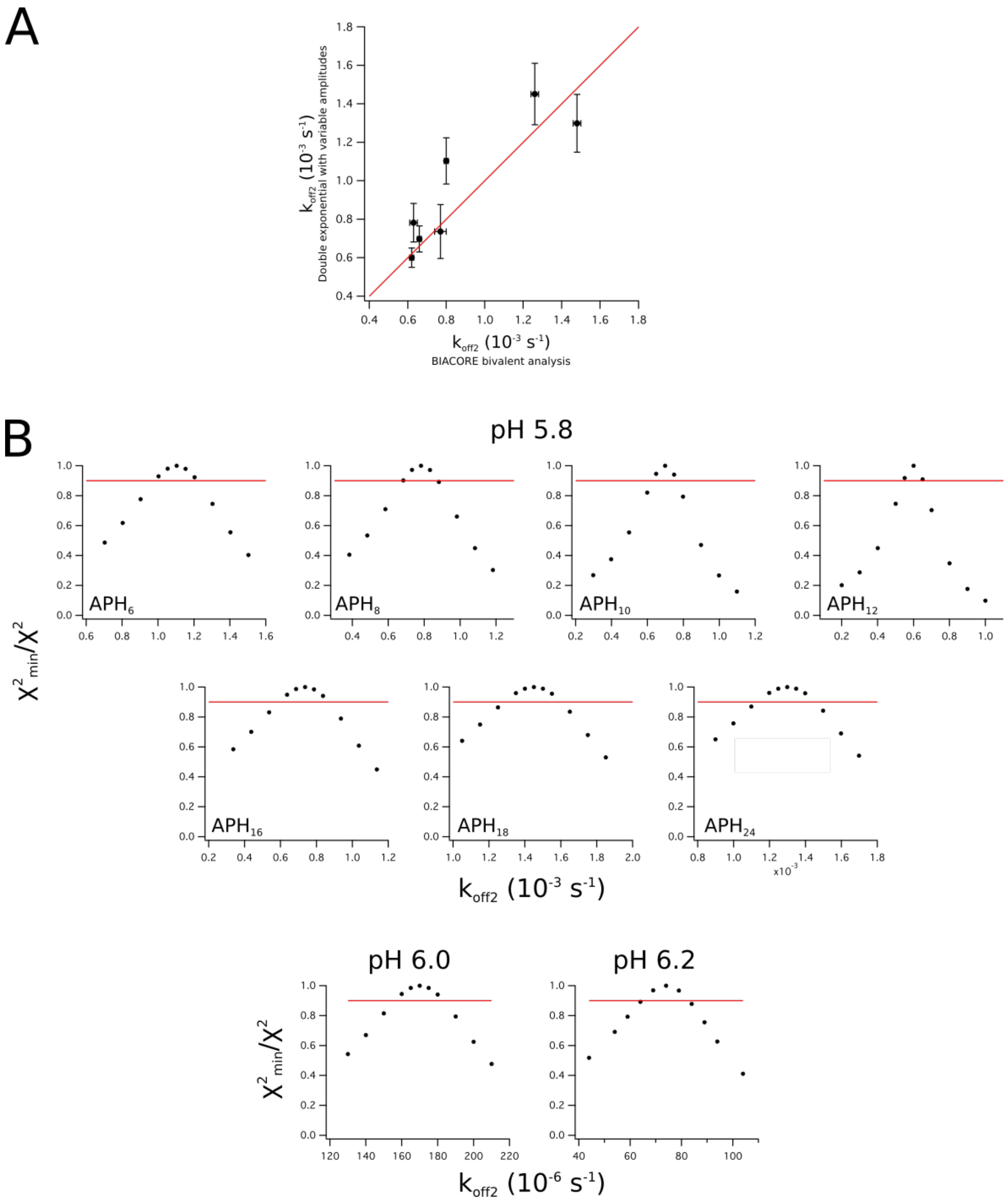


**Figure S7**. Confidence analysis of dissociation rate constants. A) The SPR data were either analysed by fitting the full data set to bivalent model in the BIACORE Evaluation software, resulting in adequate fits. We also fit the dissociation phase to a double exponential, where the amplitudes vary freely but the two rate constants are fitted globally, which improves the fit. B) To evaluate the confidence interval of the fitted rate constants, we evaluated the $\chi^2$-curves using the bivalent fit by locking $k_{off2}$ to values around the fitted value and fitting the remaining parameters. All the fitted parameters are located in well-defined $\chi^2$ minima, suggesting they are well determined by the data. Built-in error estimations typically underestimate the error associated with co-variance between fitted variables, and therefore we determined the confidence interval from a threshold of $\chi^2_{min}$ /$\chi^2$ = 0.9 (red line) following the criteria defined by Johnson et al. (Ref. 40 in main manuscript)
